# SV2A is expressed in synapse subpopulations in mouse and human brain: implications for PET radiotracer studies

**DOI:** 10.1101/2024.07.15.603608

**Authors:** Theresa Wong, Zhen Qiu, Beverly Notman, Adriana Tavares, Colin Smith, Seth G.N. Grant

## Abstract

Synapse pathology is a feature of most brain diseases and there is a pressing need to monitor the onset and progression of this pathology using brain imaging in living patients. A major step toward this goal has been the development of small-molecule radiotracers that bind to synaptic vesicle glycoprotein 2A (SV2A) for use in positron emission tomography (PET). Changes in SV2A radiotracer binding in PET are widely interpreted to report differences in the density of all synapses throughout brain regions. Here, we analyse the expression of SV2A at single-synapse resolution across regions of adult mouse and human brain. We find that SV2A is expressed in fewer than 50% of excitatory and inhibitory synapses and that the density of SV2A-positive synapses differs between brain regions. Furthermore, individual synapses differ in their amounts of SV2A. These findings have important implications for the interpretation of PET imaging studies in a clinical setting and point to the need for a detailed understanding of SV2A synaptome architecture in both healthy brain and disease cases where PET imaging is being applied.

## Introduction

Synapses - the points of communication between neurons - are present in vast numbers throughout the brain. The proteome of mammalian brain synapses comprises thousands of protein types, and mutations in their cognate genes are associated with over 130 brain disorders.^1^ These encompass a wide range of neurological and psychiatric disorders affecting individuals across the lifespan. Synapse pathology also arises from exogenous injury including stroke, trauma, inflammation, drug effects and chemical toxicity. Damage to synapses has major consequences for all aspects of brain function and commonly results in cognitive, mood, motoric and behavioural disorders. It is therefore of great importance to develop methods for monitoring synapse pathology in living humans.

Until recently, it was thought that there were comparatively few varieties of synapses in the mammalian brain. The principal classes are defined by their neurotransmitters and include excitatory (glutamatergic) and inhibitory (GABAergic), and several major classes of modulatory neurotransmitters releasing acetylcholine, dopamine, serotonin and norepinephrine, among others. However, molecular analysis has revealed an astonishing diversity of excitatory synapses arising from the differential expression of their constituent proteins^2,3^. These molecular ‘types’ of excitatory synapses, which differ in their protein composition, can be further classified into ‘subtypes’ by virtue of differences in the levels of expression, synaptic punctum morphology and nanoarchitecture.^2,4^ Importantly, these excitatory synapse types and subtypes are differentially, and characteristically, distributed such that each brain region has a compositional ‘signature’. The diversity of types and subtypes of excitatory synapses is referred to as the synaptome and their spatial distribution as the synaptome architecture^2^.

The transmembrane protein SV2A is found on synaptic vesicles in the presynaptic terminal of mammalian synapses^5^. The development of SV2A radioligands for PET imaging has garnered considerable excitement as a tool for detecting changes in synapses in humans, non-human primates and rodents^6–9^. There are now over 100 clinical studies using SV2A PET radiotracers in a wide range of disorders, including neurodegeneration (e.g. Alzheimer’s disease, frontotemporal dementia, supranuclear palsy), psychiatric disorders (e.g. schizophrenia), epilepsy and stroke.

Many publications on SV2A PET radiotracers refer to SV2A as being “ubiquitously expressed” or “ubiquitously and homogeneously expressed” in synaptic vesicles, synapses, neurons and regions of the brain^5,10–13^. As such, SV2A might be assumed a general reporter of changes in synapse density in pathology. In the present study we have employed ‘synaptome mapping’, a powerful technique that uncovers protein expression at single-synapse resolution and allows the measurement of synapse molecular diversity on a whole-brain scale. Synaptome mapping provides a first opportunity for a systematic and comprehensive examination of whether SV2A is truly ubiquitously or homogenously expressed at the level of individual excitatory and inhibitory synapses throughout regions of the mouse and human brain. The findings have important implications for the interpretation of SV2A PET imaging studies in a clinical setting and highlight the need for studies of synaptome architecture in brain disorders where SV2A imaging is being applied.

## Materials and methods

### Preparation of mouse brain tissue

All experiments were conducted in accordance with the Animal (Scientific Procedures) Act 1986, approved by British Home Office and performed by personal licence holders.

Male and female C57BL/6 adult mice (3-6 months of age) were housed at a controlled temperature with ad libitum access to food and water. Animals were anaesthetised by intraperitoneal injection of 0.1 ml pentobarbital sodium BP 20% (Dolethal, Vetoquinol UK) and transcardially perfused with 10 ml phosphate buffered saline (PBS, Oxoid), followed by 10 ml 4% paraformaldehyde (Alfa Aesar 16%, diluted 1:4 in PBS). The brains were dissected and immediately post-fixed in same fixative for 3-4 hours at 4°C. Fixed samples were cryoprotected by incubating in 5 ml 30% sucrose (VWR International) in PBS for a minimum of 48 hours at 4°C or until they sank. Brains were frozen in Optimal Cutting Temperature (OCT, CellPath) using a beaker containing isopentane (VWR Chemicals) submerged into liquid nitrogen and stored at -80°C.

### Sectioning of mouse brain tissue

Parasagittal and coronal tissue sections (18 μm thickness) were cut using a Leica CM3050S cryostat at -20°C. PBS was used as adhesive to collect tissue onto SuperFrost Plus slides (Thermo Scientific, J1800AMNZ). Slides were left to dry overnight at room temperature and stored at -20°C until use.

### Immunolabelling of synaptic proteins in mouse brain tissue

Frozen mouse brain sections were defrosted to room temperature and rinsed with Tris buffer solution (1X TBS pH 7.4, Sigma) for 5 minutes before addition of blocking solution containing 5% bovine serum albumin (BSA, Sigma), 0.5% Triton X-100 (Sigma) in 1X TBS. Slides were incubated at room temperature for 2 hours in 100 µl blocking solution prior to overnight incubation at 4°C in 100 µl primary antibody (Supplementary Table 1) and diluent solution (3% BSA, 0.2% Triton X-100 in 1X TBS). Sections were then washed three times for 10 minutes each before secondary antibody incubation (Supplementary Table 1). The washing step was repeated, followed by mounting with a glass coverslip (VWR, 1.5 thickness, 18 mm diameter) using Mowiol mounting reagent (Sigma-Aldrich) with 1,4-diazabicyclo[2.2.2]octane (DABCO). Slides were left to curate overnight in a dry staining chamber before imaging.

We employed different approaches to determine optimal immunofluorescence labelling conditions. First, we utilized C57BL/6J mouse brain tissue, which involved imaging tissue after labelling with or without primary and secondary antibodies (Supplementary Fig. 1). The SV2A antibodies produced strong punctate staining when labelled with secondary antibodies (Supplementary Fig. 1A), but signal was absent when either the primary (Supplementary Fig. 1B) or secondary antibodies were omitted. Second, we controlled for the blocking step, incubation times of antibodies and imaging parameters and derived optimal conditions based on visual inspection and signal-to-noise ratio (SNR) analysis. Lastly, to test that the SV2A-positive puncta observed (Supplementary Fig. 1C) were presynaptic terminals, we immunolabelled with a second antibody to the presynaptic protein^14^ synapsin 1 (Supplementary Fig. 1D) and observed colocalization in a subset of synapses (Supplementary Fig. 1E, arrow, inset).

### Preparation of human brain tissue

Brain tissue from deceased individuals was acquired from the University of Edinburgh Brain Bank (EBB), funded by the Medical Research Council UK. All activities involving the use of post-mortem human brain tissue received approval from the East of Scotland Research Ethics Service (21/ES/0087). Informed written consent was secured from each participant.

Human brain tissue from three control subjects (1 male, 2 female; mean age 66 ± 4.4 years) were used in this study. Control subjects did not have a history of dementia, neurological, or psychiatric disorders at time of death. The brain tissue underwent processing in accordance with EBB departmental protocols ^15^. In summary, 6 μm formalin-fixed paraffin-embedded (FFPE) tissue sections were stained with haematoxylin and eosin for routine neuroanatomical and neuropathological assessments. Examination of the brain tissue conducted by expert neuropathologist revealed minimal neuropathological changes in these control brains. Demographic and clinical characteristics are summarised in Supplementary Table 2.

During post-mortem examination, brains were macroscopically dissected into 1 cm thick coronal slices and sampled following the established EBB protocol^15^. Tissue blocks sourced from inferior temporal gyrus of the neocortex, defined as Brodmann area (BA) 20/21, were fixed in 10% formaldehyde (Genta Medical, UN2209) for 24-72 hours before tissue processing and paraffin wax embedding. Selection of anatomical region was influenced by the availability of blocks for the three control cases. Experienced pathologists conducted the macroscopic dissection of tissue blocks, closely adhering to topographical brain anatomy through the examination of sulci and gyri patterns, with assistance of a human brain atlas^16^. Adjacent Nissl-stained sections were also examined to confirm the integrity of the region of interest through assessment of distinctive cytoarchitectural features. A circular 6 mm biopsy corer (World Precision Instruments) was used to isolate smaller regions of interest from several different tissue blocks and re-embedded into a new block.

### Sectioning of human brain tissue

Coronal tissue sections (8 μm thickness) were cut using a Leica RM2125 RTS microtome at room temperature. Sections were placed into a Paraffin Section Flotation Bath (Electrothermal, MH8517) before being collected onto SuperFrost Plus slides. Slides were left to dry in an incubator (GenLab, MINO/18/SS) overnight at 40°C and stored at room temperature until use.

### Immunolabelling of synaptic proteins in human brain tissue

Human brain sections were dewaxed in two 3-minute xylene washes (Genta Medical, UN1307) and dehydrated by three 3-minute graded alcohol washes, comprising two 74°OP (IMS99%) alcohol washes and one 90% alcohol wash. To remove any formalin sedimentation, slides were washed in picric acid (TCS Biosciences, HS660-500) for 15 minutes followed by a final wash in lukewarm running water for 15 minutes. Antigen retrieval techniques were employed to improve cell surface staining, unmask antigen epitopes, and minimise non-specific background staining. The slides were then placed in 250 ml freshly made antigen retrieval solution (0.1 M sodium citrate buffer pH 6.0, Fisher Scientific) and pressure cooked (Biocare Medical, Decloaking Chamber NxGen) at 110°C for 15 minutes. Once cooled in water, human sections followed the same immunolabelling procedure as mouse protocol except for incubation time of 1 hour in blocking solution. The antibodies used to stain human brain tissue sections and their labelling conditions are described in Supplementary Table 1.

### Image acquisition

A high-resolution confocal spinning disk microscope (Nikon ECLIPSE Ti2) was used to image individual synaptic puncta in immunostained tissue (100X, NA=1.45). Obtained images measured 942 x 920 pixels in 16-bit depth with each pixel having 68 nm x 68 nm dimensions. Images were obtained through mosaic tiling whereby adjacent tiles are scanned without overlap using NIS-Elements (Nikon) software and saved in 12-bit tiff format.

### Image analysis

The individual tiles were stitched together to create a montage image using an in-house developed MATLAB script. A TrackMate plugin from Fiji software was then utlilised to detect and quantify synaptic puncta from immunofluorescence images. After manually establishing the best parameters (punctum radius; mean quality threshold; threshold) for puncta detection of each antibody marker, they were used for batch processing of puncta detection in the MATLAB implementation of the TrackMate algorithm codified in-house, whereby puncta from each channel were automatically detected and their centroid coordinates and intensity exported in text files.

Subsequently, they were imported in Python scripts developed in-house to calculate puncta density and average intensity for each brain region.

### Statistical analysis

All quantifications were collected in Excel (Microsoft Office 2016 for Windows version). Density measurements represented the number of synaptic puncta per 100 µm^2^ and intensity was quantified as arbitrary units (a.u.) representing greyscale values. All depicted values represent mean ± SEM.

## Results

We took two approaches to establishing whether SV2A is expressed ubiquitously or in subpopulations of synapses. First, we immunolabelled brain tissue sections with antibodies to SV2A and compared the density of synaptic puncta with that of synapses expressing other synaptic proteins. Second, we performed double-immunolabelling experiments using antibodies to SV2A and another synaptic protein to determine whether the two proteins colocalize in the same or distinct populations of synapses.

### Density of SV2A and other synaptic protein puncta in regions of the mouse brain

Because SV2A is a presynaptic protein, we compared its expression with four other well-characterised presynaptic proteins: synapsin 1 (SYN), synaptophysin (SYP), vesicular glutamate transporter 1 (VGLUT1) and the vesicular GABA transporter (VGAT). SYN and SYP are expressed in most presynaptic terminals, including both excitatory and inhibitory synapses^14^. To distinguish excitatory from inhibitory presynaptic terminals we used VGLUT1 and VGAT antibodies, which label each of these classes of synapse, respectively. To compare the number of SV2A presynaptic terminals with excitatory and inhibitory postsynaptic terminals, we used postsynaptic density protein 95 (PSD95) and gephyrin (GPHN) labelling, respectively. We utilised mice expressing green fluorescent protein (eGFP) fused to endogenous PSD95, which have been used extensively for mapping the diversity of excitatory synapses^2,17,18^.

We chose nine brain regions known to show high levels of synapse diversity^2^. These comprise four regions of the hippocampal formation (HPF), three regions of the neocortex, and one region each from the striatum and the thalamus. The HPF regions comprise CA1 stratum radiatum (CA1sr), CA2 stratum radiatum (CA2sr), CA3 stratum radiatum (CA3sr) and dentate gyrus polymorphic layer (DGpo). The cortical regions included layer 2/3 and 5 of the somatosensory cortex (SMSL2/3, SMSL5) and layer 4/5 of the entorhinal cortex (ENTIL4/5). The striatal caudoputamen region (STRcp) and posterior complex of the thalamus (THpo) were also examined.

Representative images from the nine mouse brain regions labelled with seven antibodies (Fig. 1) show regional variation in marker density. SV2A puncta were present at high density in CA1sr, CA2sr, CA3sr, SMSL5 and ENTIL4/5 but at low density in THpo and DGpo. Strikingly, THpo and DGpo contained very large SV2A puncta compared with the other regions. The SV2A images were similar to those for SYP, SYN and VGLUT1, but quite distinct from the VGAT puncta which were large in all brain regions. The postsynaptic markers PSD95 and GPHN showed a higher density of puncta than SV2A.

**Figure 1.**
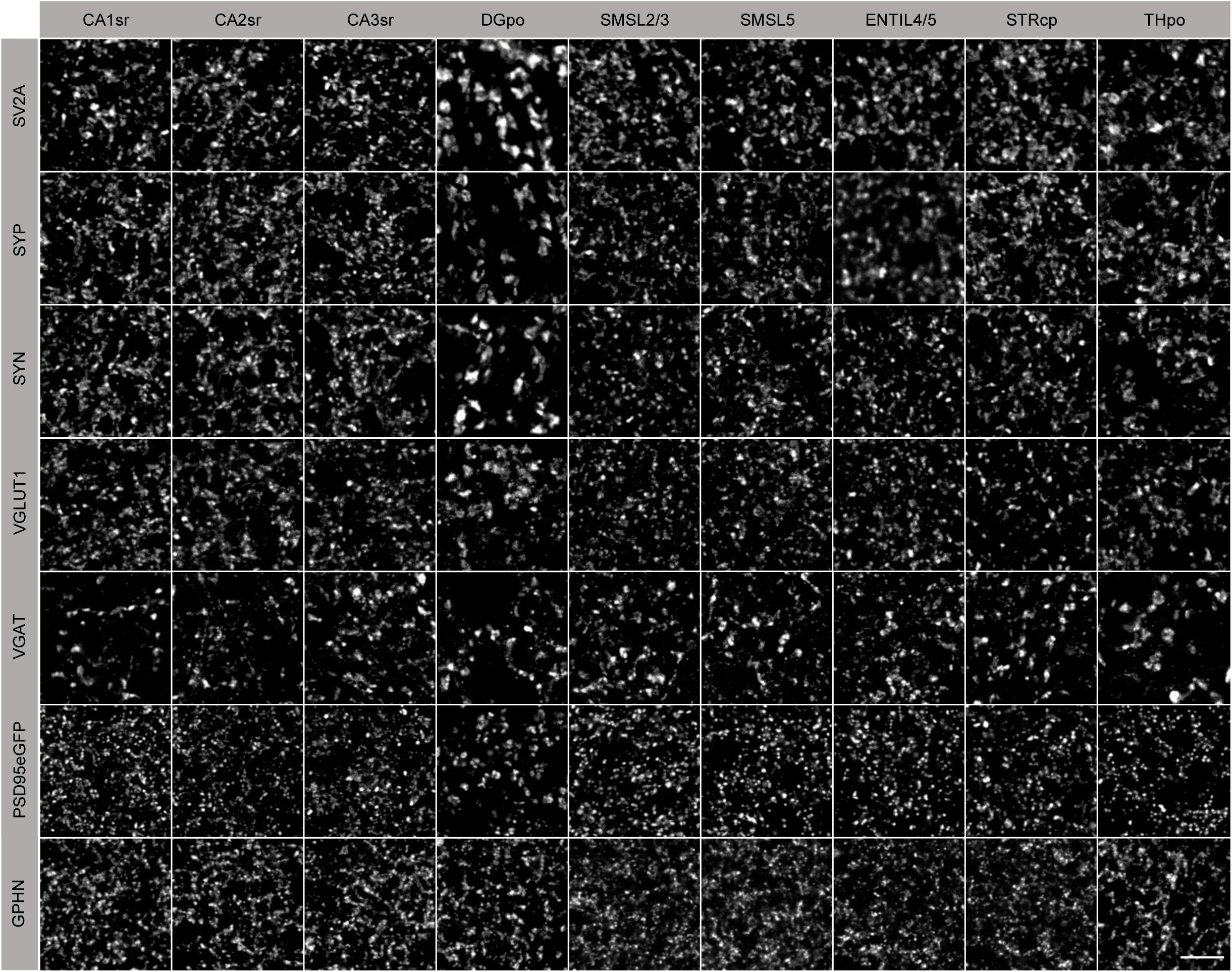
Synaptic puncta labelling in adult mouse brain regions. Nine brain regions (columns) were examined for five presynaptic (SV2A, SYP, SYN, VGLUT1, VGAT) and two postsynaptic (PSD95eGFP, GPHN) markers (rows) as described in the main text. Scale bar, 5 µm.

We quantified the density of each marker in each brain region (Supplementary Table 3) and displayed the data in a heatmap (Fig. 2A), which assists in comparing the patterns of expression between markers and regions. If SV2A were truly a universal marker for synapses, the density of SV2A should be equal to, or higher than, the combined density of the excitatory and inhibitory presynaptic markers VGLUT1 and VGAT. As shown in Supplementary Table 3, the density of SV2A puncta was similar to that of VGLUT1 and greater than that of VGAT in each brain region. However, the SV2A puncta density accounts for 60-83% of the density of both VGLUT1 and VGAT (Supplementary Table 4). This indicates that SV2A is consistently present below the combined density of VGLUT1 and VGAT, suggesting that SV2A is not expressed in all synapses, at least at the presynaptic level. In regions other than DGpo, the puncta density of the postsynaptic excitatory marker PSD95 was approximately double that of SV2A, and the puncta density of the postsynaptic inhibitory marker GPHN was higher than that of SV2A in all regions (Supplementary Table 3). SV2A puncta density accounted for only 29-40% of the density observed for both PSD95 and GPHN (Supplementary Table 5). These findings indicate that SV2A is expressed in subpopulations of excitatory and inhibitory synapses in the mouse brain.

**Figure 2.**
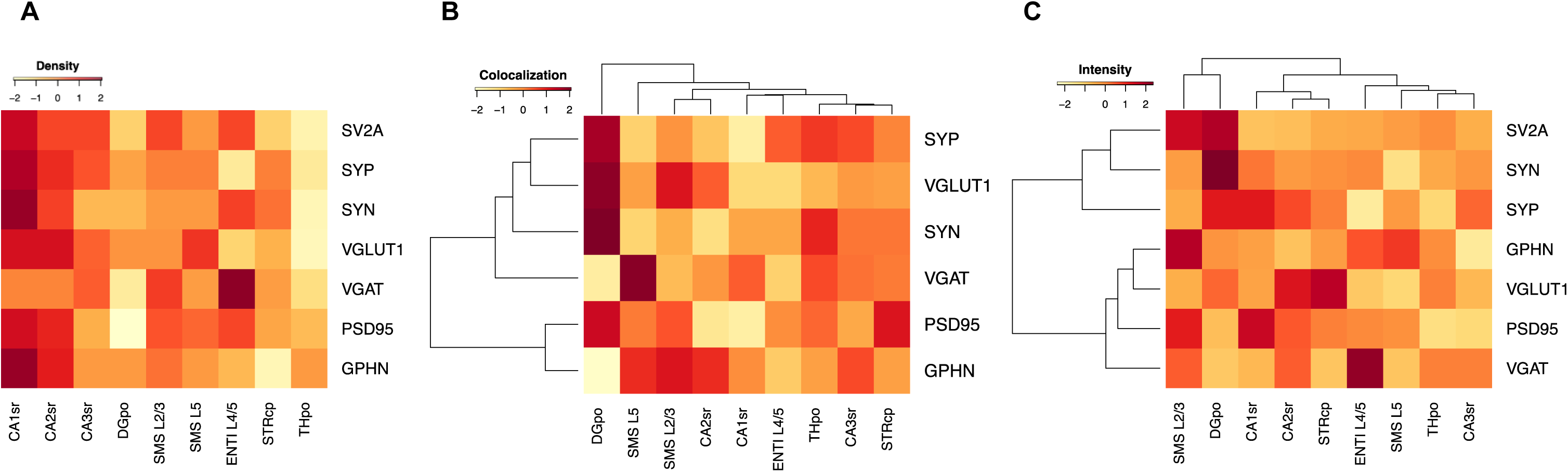
Comparison of synaptic puncta features in mouse brain regions. (**A**) Comparison of density (puncta/100 µm^2^) of synaptic puncta labelled with markers (rows) across regions of the mouse brain (columns). (**B**) Percentage colocalization of presynaptic and postsynaptic puncta with SV2A across regions of the mouse brain clustered by similarity. (**C**) Average puncta intensity (A.U.) of different synaptic proteins across regions of the mouse brain clustered by similarity. Colours are normalised values for each marker (A-C, row).

The normalised heatmap (Fig. 2A) reveals that SV2A has a similar regional distribution to SYP, SYN, VGLUT1, PSD95 and GPHN, with the highest puncta density in the CA1sr and CA2sr regions. By contrast, VGAT exhibited the highest density in ENTIL4/5.

### Colocalization of SV2A with other synaptic markers in mouse brain

The results above show that SV2A puncta density is lower than that of the sum of excitatory and inhibitory markers at both the presynaptic and postsynaptic levels. From this we would expect SV2A to be present in subsets of excitatory and inhibitory synapses. To test this hypothesis, we conducted dual-labelling experiments using SV2A with the excitatory or inhibitory presynaptic marker VGLUT1 or VGAT. Representative images (Fig. 3) clearly demonstrate the existence of synapses expressing either protein, as well as those expressing both markers. Quantification of these data shows that in the regions examined SV2A was present in 57-84% of VGLUT1 synapses and in 52-92% of VGAT synapses (Table 1). Furthermore, we investigated the juxtaposition of SV2A with the postsynaptic markers PSD95 and GPHN (Fig. 3, Table 1). In the brain regions examined, 32-49% (mean = 41%) of excitatory postsynaptic terminals and 38-45% (mean = 41%) of inhibitory postsynaptic terminals were found adjacent to presynaptic terminals positive for SV2A. Together, these findings provide strong evidence that SV2A is present in specific subsets of excitatory and inhibitory synapses.

**Figure 3.**
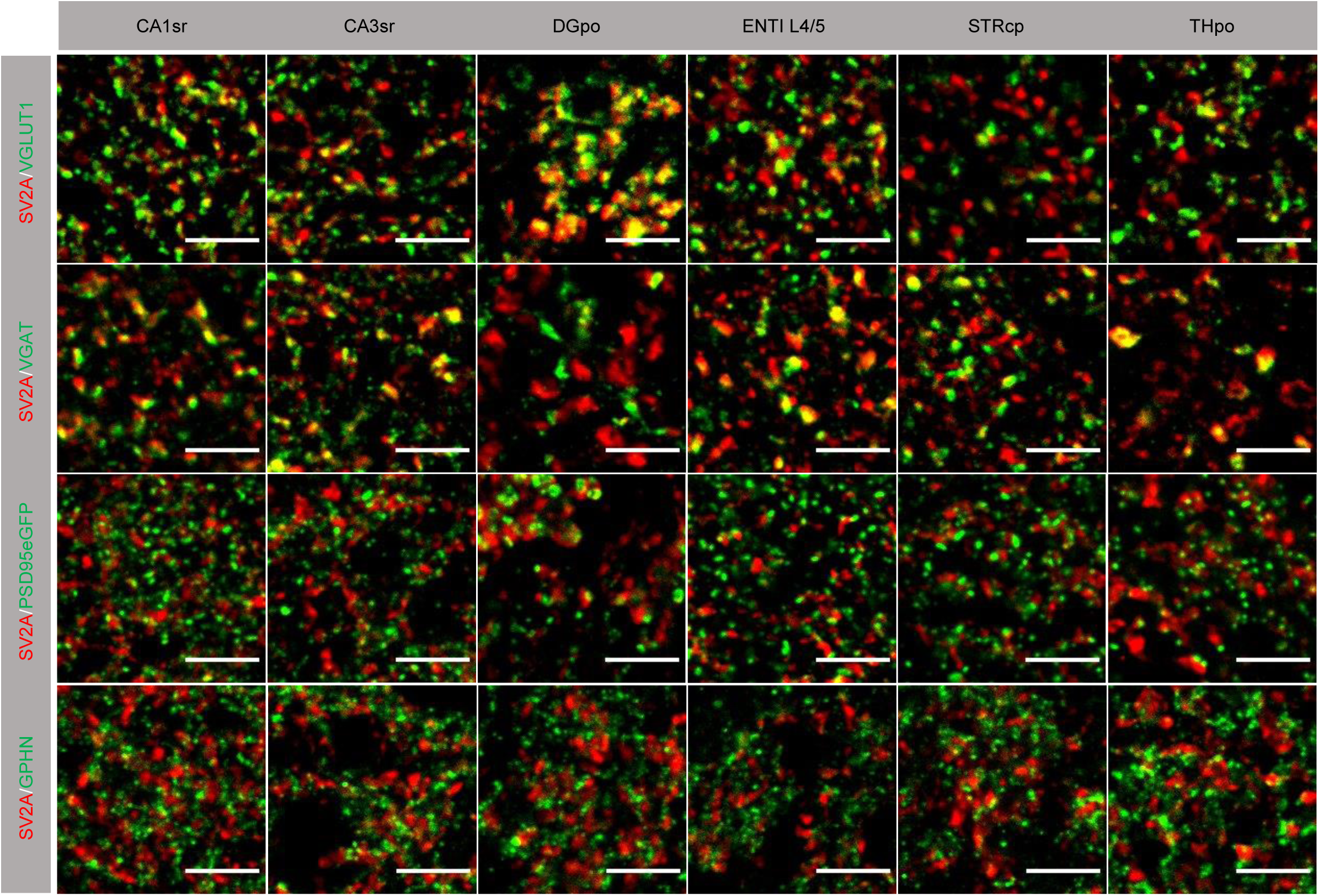
Synaptic colocalisation of SV2A with presynaptic and postsynaptic markers in adult mouse brain. Images of individual synaptic puncta labelled with SV2A (red) and excitatory (VGLUT1, PSD95eGFP) and inhibitory (VGAT, GPHN) synaptic markers (green). Yellow puncta indicate colocalization of SV2A with the other markers. Scale bars, 5 µm.

**Table 1.** Colocalization of SV2A with excitatory and inhibitory synaptic markers across regions of mouse brain. Values (mean ± SEM) represent percentage of marker puncta that contain SV2A+ puncta.

| Marker/ROI | CA1sr | CA2sr | CA3sr | DGpo | SMS L2/3 | SMS L5 | ENTIL4/5 | STRcp | THpo | Mean |
| --- | --- | --- | --- | --- | --- | --- | --- | --- | --- | --- |
| SYP | 65±1 | 70±8 | 78±5 | 85±3 | 74±1 | 69±2 | 77±7 | 75±3 | 79±4 | 75±4 |
| SYN | 57±10 | 49±11 | 62±12 | 80±6 | 56±19 | 51±19 | 57±13 | 62±13 | 69±11 | 60±13 |
| VGLUT1 | 57±2 | 70±5 | 65±3 | 84±6 | 77±6 | 64±4 | 57±6 | 64±1 | 61±8 | 67±5 |
| VGAT | 74±3 | 68±7 | 72±7 | 52±8 | 62±5 | 92±12 | 59±9 | 71±7 | 76±6 | 69±7 |
| PSD95 | 32±9 | 33±5 | 39±3 | 49±6 | 44±2 | 42±1 | 41±7 | 48±3 | 43±3 | 41±4 |
| GPHN | 41±2 | 44±3 | 43±3 | 34±1 | 45±2 | 44±1 | 38±3 | 40±6 | 40±3 | 41±3 |

We next asked if SV2A preferentially colocalizes with any particular synaptic markers in the different brain regions using a hierarchically clustered heatmap (Fig. 2B). There was no single region where SV2A exhibited high colocalization with all markers. Instead, a dichotomy was evident: within the DGpo (which contains the largest SV2A puncta) SV2A had the highest colocalization with VGLUT1 and lowest colocalization with VGAT. The opposite was observed in SMSL5. These findings indicate that SV2A does not consistently represent excitatory and inhibitory synapses in all brain regions.

### Variation in the amount of SV2A protein in individual synapses

The visual examination of synaptic puncta labelled with SV2A and other markers revealed a wide range of sizes and intensities (Fig. 1). Some brain regions, such as the DGpo, consistently contained larger and more intensely labelled presynaptic terminals. To analyse the synapse populations in each brain region in a quantitative manner, we calculated the mean puncta intensity for each marker in each region (Supplementary Table 6) and, to assist in comparing these data, we created a hierarchically clustered heatmap (Fig. 2C).

As shown in Fig. 2C, SV2A puncta exhibited highest mean intensity in SMSL2/3 and DGpo, as compared with other brain regions. Comparing the cortical layers shows that SV2A puncta have higher intensity in the superficial (SMSL2/3) than deeper (SMSL5, ENTIL4/5) layers; a distribution pattern that was not observed for any of the other markers (Fig. 2C). SYN and SYP clustered with SV2A, whereas the other markers formed a separate cluster. It is worth noting that like SV2A, SYN and SYP are markers that do not distinguish between inhibitory and excitatory synapses. By contrast, markers specific to either excitatory or inhibitory synapses formed a distinct cluster, as shown in Supplementary Table 6 and in Fig. 2C.

### SV2A is expressed in subsets of human brain synapses

To investigate whether SV2A is also expressed in subsets of synapses in the human brain, sections from the inferior temporal gyrus (BA20/21) of the neocortex (layers 1 to 6) were immunolabelled for SV2A, SYP, PSD95 or GPHN (Fig. 4). Punctate staining was observed for all markers, with SV2A and SYP labelling larger puncta than PSD95 and GPHN. Variation in puncta shape, size and intensity was observed across cortical layers.

**Figure 4.**
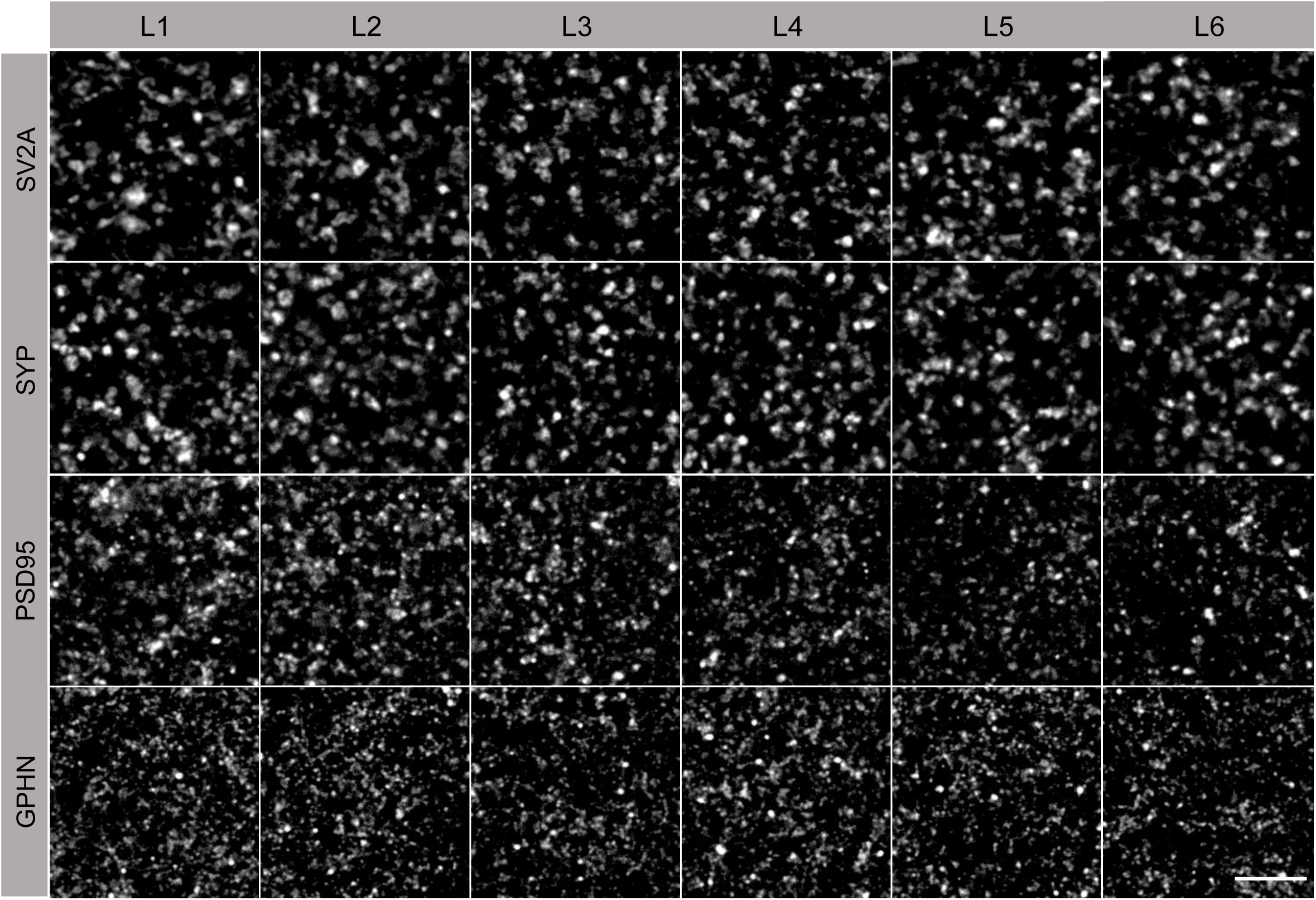
Antibody labelling of synapses in human temporal cortex. Individual synaptic puncta in the six layers (L1-L6) of the human temporal cortex labelled with presynaptic (SV2A, SYP) and postsynaptic (PSD95, GPHN) markers. Scale bar, 5 µm.

We first compared the density of the four markers in each of the six layers. The density of SV2A and SYP puncta was approximately half that of PSD95 or GPHN in each layer (Supplementary Table 7). The density of SV2A and SYP puncta was similar in all layers, in contrast to PSD95 and GPHN which showed a gradient, with the higher puncta density in superficial layers and lower density in deeper layers (Supplementary Table 7). Comparing the density of SV2A puncta with the total (sum of PSD95 and GPHN) density revealed that SV2A accounted for 23-27% across layers and 25% mean overall (Supplementary Table 8). These data indicate that SV2A labels ∼25% of synapses in the human neocortex.

We next examined the colocalization of SV2A with SYP, PSD95 or GPHN (Fig. 5, Table 2). Although SV2A and SYP have a similar density in each cortical layer (Supplementary Table 7), overall across all layers 67% of SYP synapses express SV2A (Table 2). Only 30% of PSD95 synapses and 23% of GPHN synapses were labelled with SV2A (Table 2).

**Figure 5.**
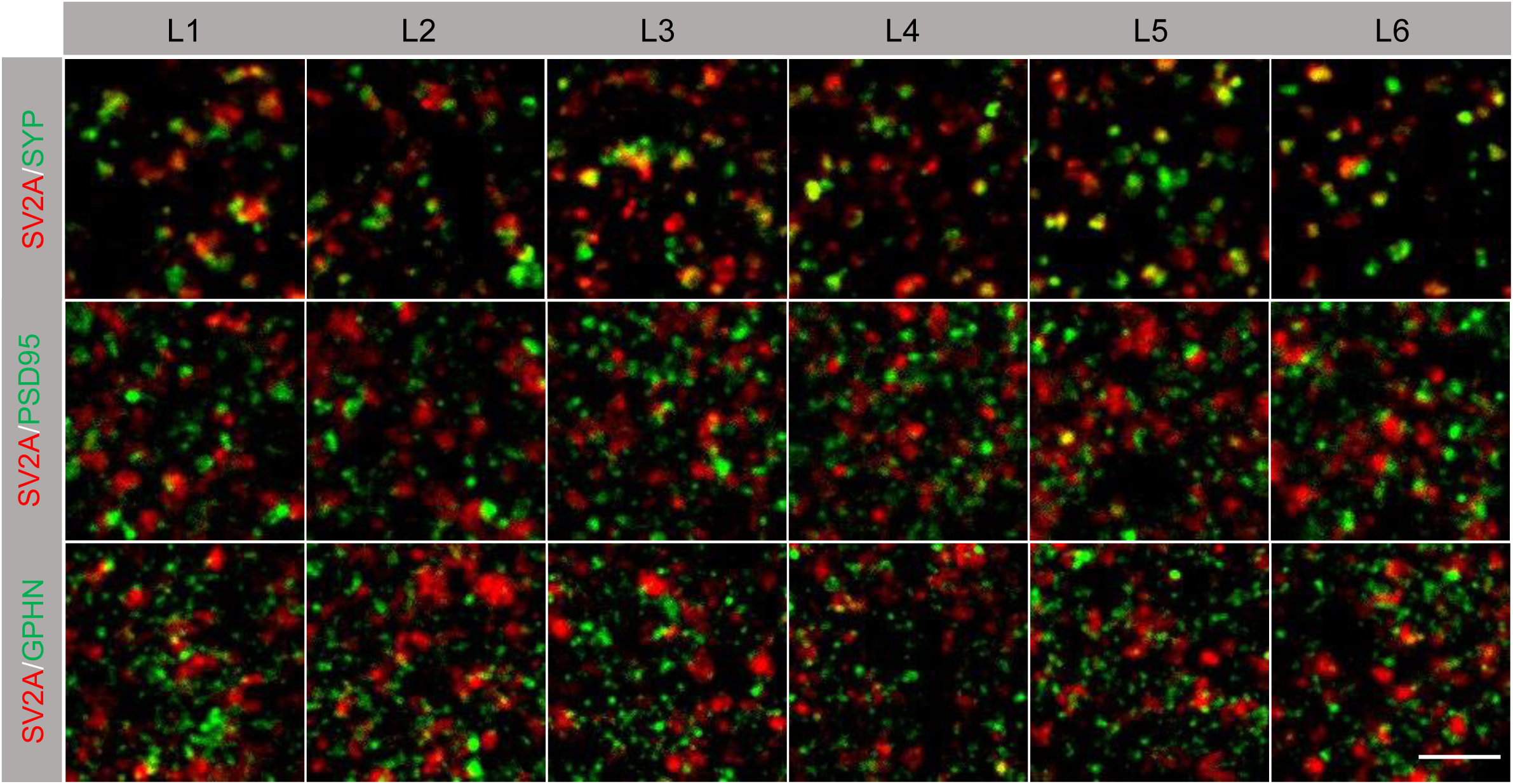
Synaptic colocalisation of SV2A with presynaptic and postsynaptic markers in human brain. Images of individual synaptic puncta in layers (L1-L6) of human temporal cortex labelled with SV2A (red) and presynaptic marker SYP, postsynaptic excitatory synapse marker PSD95, and postsynaptic inhibitory marker GPHN (all green). In SV2A/SYP labelling (top row), yellow indicates colocalization of the two markers. In SV2A/PSD95 (middle row) and SV2A/GPHN (bottom row) labelling, the juxtaposition of red and green puncta indicates synaptic colocalization. Scale bar, 5 µm.

**Table 2.**
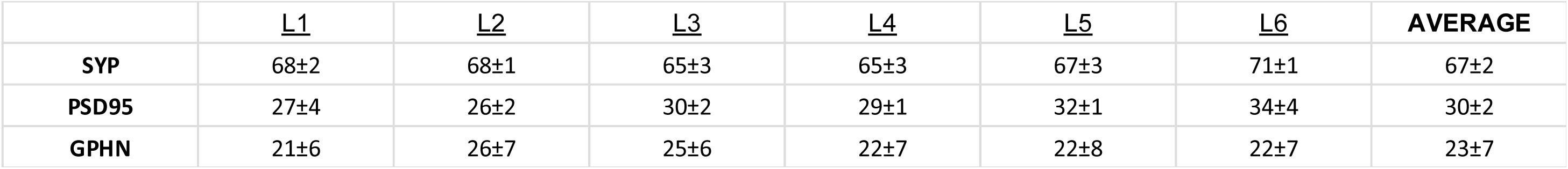
Percentage of SV2A colocalization with other synaptic markers in layers of human temporal cortex. Values (mean ± SEM) represent percentage of marker puncta that contain SV2A+ puncta.

|  | <u>L1</u> | <u>L2</u> | <u>L3</u> | <u>L4</u> | <u>L5</u> | <u>L6</u> | **AVERAGE** |
| --- | --- | --- | --- | --- | --- | --- | --- |
| **SYP** | 68±2 | 68±1 | 65±3 | 65±3 | 67±3 | 71±1 | 67±2 |
| **PSD95** | 27±4 | 26±2 | 30±2 | 29±1 | 32±1 | 34±4 | 30±2 |
| **GPHN** | 21±6 | 26±7 | 25±6 | 22±7 | 22±8 | 22±7 | 23±7 |

Finally, we examined the amount of each marker in synapses by measuring the mean intensity of individual puncta in each brain region (Fig. 6, Supplementary Table 9). As shown in the heatmap, layer 1 contains synapses with the most intense puncta. SV2A intensity was lowest in layer 4, SYP in layer 2, PSD95 and GPHN in layer 6.

**Figure 6.**
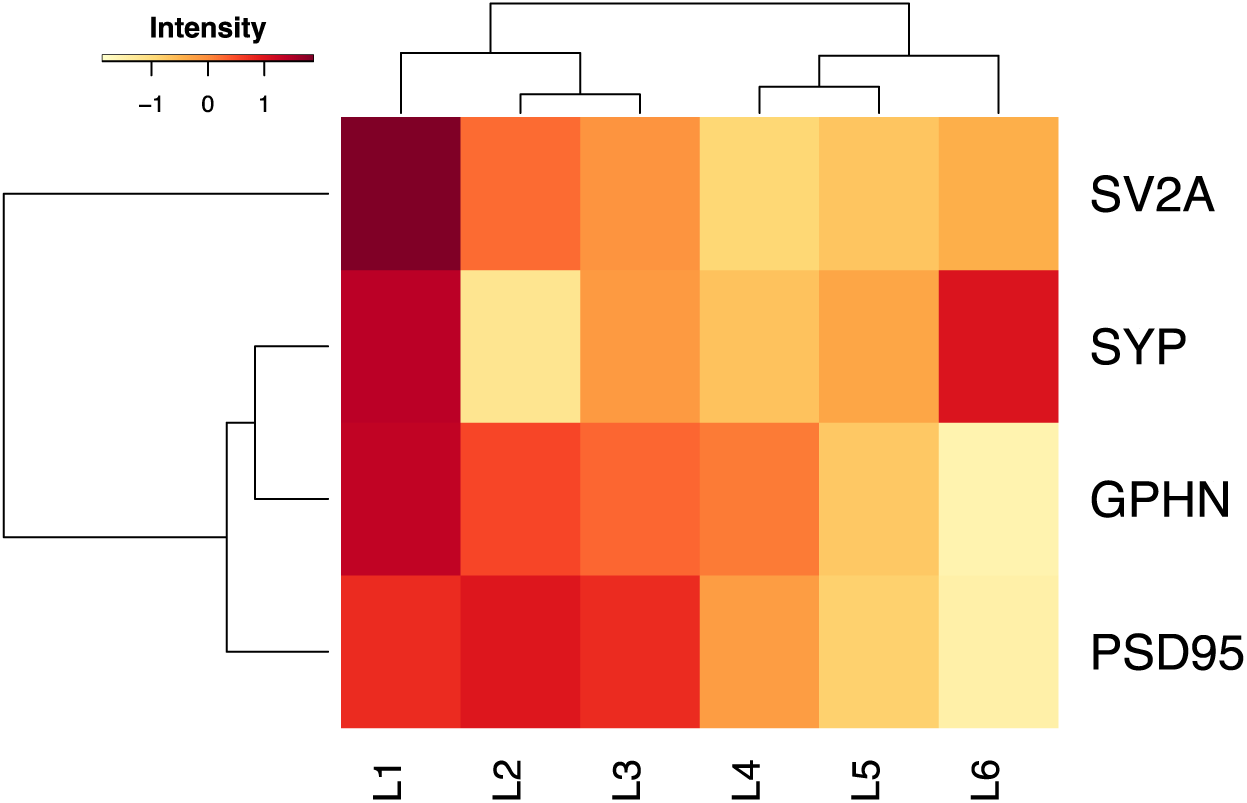
Comparison of intensity of presynaptic and postsynaptic puncta across layers of human temporal cortex. Average puncta intensity (A.U.) of four synaptic proteins across layers (L1-L6) of the human temporal cortex clustered by similarity. Colours are normalised values for each marker (row).

### Differences in SV2A between human and mouse brain

We observed some differences between the mouse and human brain. Comparing L2 in the mouse cortex (SMSL2/3) with L2 in human temporal cortex, the density of SV2A, SYP, PSD95 and GPHN puncta was consistently higher in human (Supplementary Tables 3, 7). The percentage colocalization of pairs of markers also differed between mouse cortex (SMSL2/3) with L2 in human temporal cortex (Tables 1, 2). Averaging all regions, the mean percentage colocalization of SV2A with either PSD95 or GPHN for all regions examined was higher in the mouse (PSD95: mouse 41%, human, 30%; GPHN: mouse 41%, human 23%) (Tables 1, 2). Although this comparison does not compare the exact same regions of the two species, the current findings nonetheless suggest that there are differences in the composition of the synaptome between mouse and human brain regions, likely due to evolutionary divergence^19^.

## Discussion

The presynaptic protein SV2A has gained clinical importance because it is the target of PET radiotracers used to measure synapse density in the brain of living patients. It is also the target of anti-epileptic drugs. Understanding how widely SV2A is expressed in the brain, and whether it reports all or only subpopulations of synapses, are clearly of relevance to its clinical applications. Using high-resolution, wide-scale spinning disk confocal microscopy we have quantified the number of synapses expressing SV2A, and the levels of expression in individual synapses, in conjunction with other presynaptic and postsynaptic proteins in mouse and human brain. These studies reveal that SV2A is expressed in subpopulations of excitatory and inhibitory synapses, the density of which varies across mammalian brain regions, and that there is high diversity among individual SV2A-positive synapses.

In the temporal cortex of adult humans, we found that ∼25% of all synaptic puncta are SV2A positive. Only 30% of excitatory and 23% of inhibitory synapses expressing PSD95 and GPHN, respectively, colocalized with SV2A. These fractions varied across the layers of the cortex. Approximately 67% of SYP-positive synapses expressed SV2A across cortical layers. We observed similar results in the mouse brain. In our analysis of cortical, hippocampal, striatal and thalamic regions, SV2A was expressed in subpopulations of excitatory and inhibitory synapses. Overall, SV2A puncta accounted for 29-40% of PSD95 and GPHN puncta in different brain regions. Examination of SV2A colocalization with presynaptic markers in different regions showed that 57-84% of VGLUT1 synapses and 52-92% of VGAT synapses expressed SV2A. At the postsynaptic level, SV2A colocalized with 32-49% of excitatory and 38-45% of inhibitory postsynaptic terminals. Similar to human brain, 65-85% of SYP-positive synapses also expressed SV2A. Together, these findings show that in many regions of the mammalian brain, SV2A radiotracers will likely detect fewer than 50% of synapses. Moreover, this fraction varies between brain regions and between excitatory and inhibitory synapses. We also observed considerable heterogeneity of SV2A-positive synapses with respect to their size and protein content.

That SV2A exhibits synapse diversity is fully in line with previous investigations into the synaptome of other synaptic proteins. Studies in the mouse brain show that a diversity of synapse types can be distinguished by their molecular composition (synapse proteomes) and that these may be further divided into subtypes according to punctum size, shape, nanoarchitecture, protein lifetime and level of expression^2,4,17,18^. Together, these synapse types and subtypes comprise the synaptome of the brain.

### SV2A PET radiotracers as reporters of synapse density

SV2A is commonly described as being expressed in all synapses at homogenous levels and in all brain regions. To understand how this view arose and evolved in the PET literature, we identified over 60 studies that refer to its expression as ubiquitous. The vast majority of this literature refers to two papers^6,13^. The earliest is from Bajjelieh and colleagues, who used mRNA in situ hybridisation on rat brain tissue sections (imaged at low resolution) and describe “ubiquitous expression” in the context of brain regions and suggest that their results are “consistent” with expression of the gene in all neurons. They do not show SV2A protein expression at single-synapse resolution. The latter paper from Finnema and colleagues refers to SV2A as “ubiquitously and homogeneously located in synapses across the brain” and cites the Bajjelieh study. It is notable that in Figure 1K of their paper they show immunofluorescent labelling of SV2A and SYP synaptic puncta in a region of grey matter and it is clear from this image that the two proteins have differential and overlapping expression, consistent with our findings. To our knowledge, studies that directly correlate brain regional or pathological single-synapse resolution quantification of synapse density with PET binding data remain absent from the published literature.

### Implications of synapse diversity for the clinical use of PET imaging

In considering the impact of our findings on the interpretation of PET imaging data it is useful to contrast two models: the ‘standard model’ where SV2A is expressed ubiquitously and homogeneously, and the ‘synaptome model’ of the present study in which SV2A is expressed heterogeneously in 25% of synapses. Our findings point to five relevant factors:

1. SV2A radiotracers likely bind to a minority of the total number of brain synapses;
2. SV2A is present in different proportions of excitatory and inhibitory synapses, with a higher fraction of inhibitory synapses expressing SV2A;
3. The fraction of all synapses expressing SV2A differs between brain regions, as does the fraction of excitatory and inhibitory synapses;
4. The amount of SV2A varies between individual synapses;
5. The average amount of SV2A in the populations of synapses in brain regions differs.

The design of most clinical studies with SV2A PET imaging involves a comparison of a set of cases (e.g. patients with a brain disorder) compared with controls. We will consider a hypothetical study that reports a 20% reduction in SV2A PET radiotracer binding in two brain regions (A and B) of patients. First, the SV2A radiotracer tells us nothing about the 75% of brain synapses that do not express SV2A. Instead, the synaptome model interprets the 20% reduction in signal as a 5% reduction in the total number of synapses, as only 25% of synapses express SV2A. Furthermore, the synaptome model tells us that this might differentially impact excitatory versus inhibitory synapses, and region A versus region B (i.e. the extent of pathology may differ between those regions), especially if synapses in these two regions express different levels of SV2A.

Factors 4 and 5 are important because the capacity to detect the loss of a synapse is affected by the amount of protein in that synapse. PET radiotracers occupy a small fraction of available binding sites, hence synapses that express small amounts of protein will be more likely to escape detection in PET imaging. For example, if region A is composed of large SV2A synapses with high-intensity expression and region B has small low-intensity synapses, then a 20% reduction in binding in region B will mean a greater loss of synapses.

It is important to recognise that disease processes may interfere with any of the steps (expression of the gene, translation of the protein in soma or axon, transport of the protein to the synapse, assembly into the synaptic vesicle) that determine the amount of SV2A in a synapse. For example, Pazalar and colleagues found that the level of *SV2A* mRNA in single neurons isolated from patients with epilepsy was reduced in many types of neurons^20^. Importantly, the level of SV2A mRNA and/or protein has no impact on the number of synapses, as mice lacking SV2A show no loss of synapse density or change in synapse morphology^21^. This finding, together with our present study, emphasises that a change in SV2A radiotracer binding does not necessarily imply a change in synapse density as it could equally reflect a (pathological) change in the level of the protein.

### Recommendations for interpretation of SV2A PET imaging studies

We argue that changes in SV2A PET radiotracer binding might not reflect a change in synapse density, or may greatly overestimate it, or may fail to detect major losses of SV2A-negative synapses. Consequently, in studies that employ PET radiotracers to study human brain disease, we caution against the conclusion that altered binding be interpreted as a direct reporter of pathological change in synapse density. Instead, changes in SV2A PET radiotracer binding should be interpreted as changes in SV2A-positive synapses. Similar reservations may impact the interpretation of SV2A radiotracer binding studies undertaken for the normal brain. A comprehensive brain-wide, single-synapse resolution examination of the distribution of SV2A is called for, incorporating further synaptic markers that enable any change in synaptic populations to be quantified with confidence. Synaptome mapping technology is ideally suited to this objective as it has been demonstrated that brain-wide synaptome analysis of PSD95-positive synapses is possible in human brain^22^.

Such a detailed knowledge of the synaptome architecture of SV2A across the brain, and lifespan, could greatly enhance the discovery value of SV2A PET imaging. For example, in Alzheimer’s disease, which is characterised by synapse loss during the early stages, it would be informative to determine whether and how SV2A-positive synapse types and subtypes are affected at the different stages of disease progression. One of the key principles that has emerged from synaptomic theory is that synaptic proteins that are encoded by disease-risk genes are expressed in subpopulations of synapses^23,24^. Indeed, *SV2A* is recognised as a schizophrenia risk gene^25^ and thus SV2A-positive synapses are candidates for the underlying pathology of this disorder. Thus, it would be valuable to characterise expression of other proteins in SV2A-positive synapses as this might indicate that SV2A PET studies could be particularly useful in these disorders. It is also noteworthy that SV2A is the target of a highly effective anti-epileptic drug and the identification of the SV2A-positive subpopulations of synapses may inform on its mechanism of action and lead to new therapies. In conclusion, detailed synaptome analysis of SV2A on the human brain in health and disease will greatly enhance the value of SV2A PET radiotracers and advance their application in clinical settings.

## Acknowledgements

Laboratory management, E. Sigfridsson. Edinburgh Brain Bank, C. McKenzie. Discussions, E. Bulovaite. Manuscript editing, C. Davey. For the purpose of open access, the author has applied a Creative Commons Attribution (CC-BY) licence to any Author Accepted Manuscript version arising from this submission.

## Author contributions

Laboratory experiments: TW, BN. Image analysis: TW, ZQ. Human brain samples: CS. Supervision: AT, CS, SGNG. Writing: TW, SGNG.

## Funding

This research was funded by the Wellcome Trust (221295/Z/20/Z, 218293/Z/19/Z).

## Competing interests

The authors report no competing interests.

## Data availability

Data is available on request.

## Supplementary material

Supplementary material is available at *Brain* online.

---

**Supplementary Figure 1.**
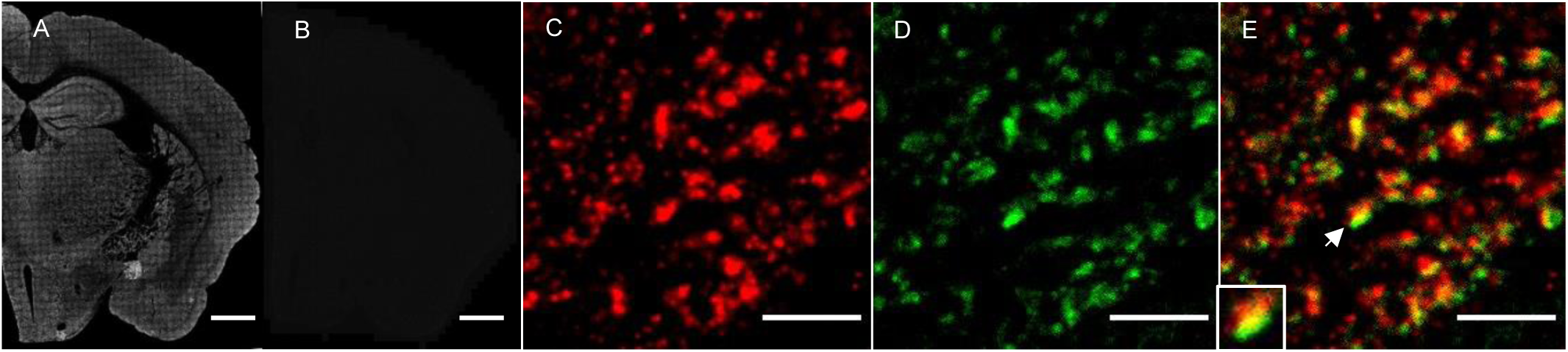
SV2A immunofluorescence optimisation staining methods on adult mouse brain. (**A**,**B**) Low-magnification image of coronal section of mouse brain stained with anti-SV2A rabbit polyclonal antibody and secondary antibody labelled with Alexa Fluor 647 (A). An adjacent brain section was stained without primary or secondary antibody as negative control (B). Scale bars, 1 mm. (**C-E**) High-magnification image (100x objective) of CA1sr regions of mouse hippocampal formation labelled with antibody to SV2A (C), SYN (D) and merged (E). Scale bars, 5 µm. An example of marker colocalisation (arrow) is enlarged in the inset.

**Supplementary Table 1.** Primary and secondary antibodies used in this study. AF488, Alexa Fluor 488; AF647, Alexa Fluor 647; CY5, Cyanine5.

| Antibody | Species | Source | Catalogue Number | Dilution | Concentration | Tested in | RRID |
| --- | --- | --- | --- | --- | --- | --- | --- |
| SV2A | Rabbit | Abcam | ab32942 | 1:500, 1:250 | 1 mg/mL | M, H | AB_778192 |
| SYP | Mouse | Synaptic Systems | 101 111 | 1:500, 1:250 | 1 mg/mL | M, H | AB_2782972 |
| SYN | Mouse | Synaptic Systems | 106 011 | 1:500 | 1 mg/mL | M | AB_2619772 |
| VGLUT1 | Mouse | Neuromab | 75-066 | 1:500 | 1 mg/mL | M | AB_2187693 |
| VGAT | Guinea pig | Synaptic Systems | 131 005 | 1:500 | 1 mg/mL | M | AB_1106810 |
| PSD95 | Mouse | Synaptic Systems | 124 011 | 1:250 | 1 mg/mL | H | AB_10804286 |
| GPHN | Mouse | Neuromab | 75-465 | 1:500, 1:250 | 1 mg/mL | M, H | AB_2716264 |
| Anti-rabbit AF488 | Goat | Thermo Fisher Scientific | A-11008 | 1:1000, 1:500 | 2 mg/mL | M, H | AB_143165 |
| Anti-rabbit AF647 | Goat | Thermo Fisher Scientific | A-21244 | 1:1000, 1:500 | 2 mg/mL | M, H | AB_2535812 |
| Anti-mouse CY5 | Goat | Jackson ImmunoResearch Laboratories | 115-175-205 | 1:1000, 1:500 | 1.2 mg/mL | M, H | AB_2338715 |
| Anti-guinea pig AF647 | Goat | Thermo Fisher Scientific | A-21450 | 1:1000 | 2 mg/mL | M | AB_2735091 |

**Supplementary Table 2.** Demographic and clinical characteristics of human subjects. BMI, body mass index; CAT, coronary artery atherosclerosis; CIEDs, cardiac implantable electronic devices; DCM, dilated cardiomyopathy; HCL, hypercholesterolemia; HHD, hypertensive heart disease; HTN, hypertension; IHD, ischaemic heart disease; LBBB, left bundle branch block; MI, myocardial infarction; PE, pulmonary embolism; PMD, post-mortem delay; PMH, past medical history; T1D, type 1 diabetes mellitus.

| Case ID | SD014/13 | SD046/17 | SD003/14 |
| --- | --- | --- | --- |
| Age (years) | 74 | 65 | 59 |
| Gender | Female | Female | Male |
| Height (m) | 1.63 | 1.63 | 1.75 |
| Weight (kg) | 82 | 57 | 83 |
| BMI | 50.3 | 35 | 47.4 |
| Medication | No | No | No |
| Significant PMH | HTN | HTN, DCM, CIEDs, LBBB | HCL, T1D |
| Cause of death | PE | IHD, HHD | MI, CAT, T1D |
| Brain weight (g) | 1520 | 1150 | - |
| PMD (hr) | 41 | 76 | 50 |
| pH | 6.3 | 6.35 | 6.1 |

**Supplementary Table 3.** Quantification of synaptic puncta density of excitatory and inhibitory protein markers across regions the mouse brain. Measurements normalised to puncta/100 µm^2^. Mean ± SEM. ROI, region of interest.

| Marker/ROI | CA1sr | CA2sr | CA3sr | DGpo | SMS L2/3 | SMS L5 | ENTIL4/5 | STRcp | THpo | Mean |
| --- | --- | --- | --- | --- | --- | --- | --- | --- | --- | --- |
| SV2A | 19±1 | 18±1 | 18±1 | 16±1 | 18±1 | 17±1 | 18±1 | 16±1 | 15±1 | 17±1 |
| SYP | 17±3 | 15±3 | 14±3 | 12±1 | 13±1 | 13±1 | 9±6 | 13±4 | 9±4 | 13±3 |
| SYN | 17±3 | 14±3 | 11±2 | 11±1 | 12±1 | 12±1 | 14±4 | 13±4 | 8±4 | 12±2 |
| VGLUT1 | 19±1 | 19±3 | 17±3 | 16±3 | 16±2 | 18±3 | 14±1 | 15±1 | 12±1 | 16±2 |
| VGAT | 10±3 | 10±4 | 11±3 | 5±2 | 12±3 | 9±1 | 16±3 | 9±1 | 6±2 | 10±2 |
| PSD95 | 39±1 | 37±2 | 28±3 | 18±3 | 34±1 | 33±1 | 35±1 | 29±4 | 27±1 | 31±2 |
| GPHN | 27±1 | 25±1 | 22±2 | 22±1 | 23±1 | 22±1 | 21±3 | 18±2 | 22±1 | 22±1 |

**Supplementary Table 4.** Comparison of SV2A, VGLUT and VGAT puncta density in mouse brain regions. SV2A puncta density (top row), summed density of VGLUT1 and VGAT (middle row), and the percentage of SV2A puncta compared with VGLUT1 and VGAT (bottom row).

| Marker/ROI | CA1sr | CA2sr | CA3sr | DGpo | SMS L2/3 | SMS L5 | ENTIL4/5 | STRcp | THpo | Mean |
| --- | --- | --- | --- | --- | --- | --- | --- | --- | --- | --- |
| SV2A | 19 | 18 | 18 | 16 | 18 | 17 | 18 | 16 | 15 | 17 |
| VGLUT1 + VGAT | 29 | 29 | 28 | 21 | 28 | 27 | 30 | 24 | 18 | 26 |
| % SV2A | 66 | 62 | 64 | 76 | 64 | 63 | 60 | 67 | 83 | 65 |

**Supplementary Table 5.**
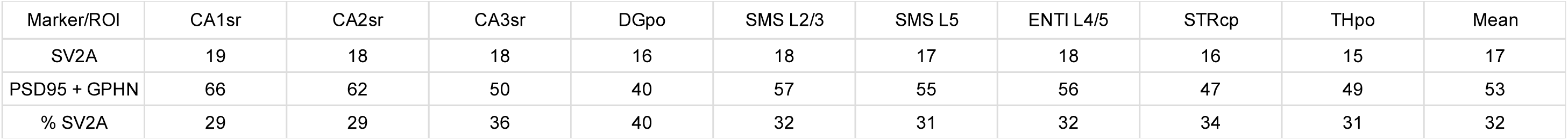
Percentage of SV2A-positive puncta compared with postsynaptic markers in mouse brain regions. SV2A puncta density (top row), summed density of PSD95 and GPHN (middle row), and the percentage of SV2A puncta compared with PSD95 and GPHN (bottom row).

**Supplementary Table 6.**
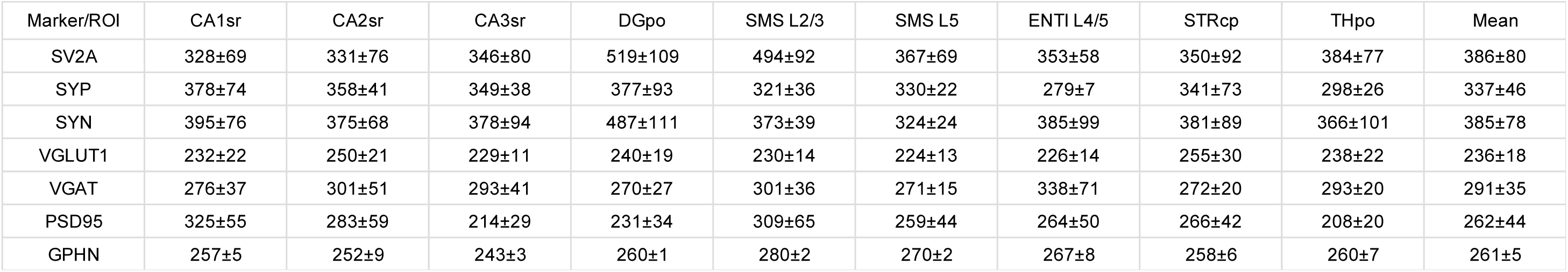
Mean intensity expression of protein markers in individual synapses in mouse brain regions. Intensity values (a.u.) shown as mean ± SEM.

| Marker/ROI | CA1sr | CA2sr | CA3sr | DGpo | SMS L2/3 | SMS L5 | ENTIL4/5 | STRcp | THpo | Mean |
| --- | --- | --- | --- | --- | --- | --- | --- | --- | --- | --- |
| SV2A | 328±69 | 331±76 | 346±80 | 519±109 | 494±92 | 367±69 | 353±58 | 350±92 | 384±77 | 386±80 |
| SYP | 378±74 | 358±41 | 349±38 | 377±93 | 321±36 | 330±22 | 279±7 | 341±73 | 298±26 | 337±46 |
| SYN | 395±76 | 375±68 | 378±94 | 487±111 | 373±39 | 324±24 | 385±99 | 381±89 | 366±101 | 385±78 |
| VGLUT1 | 232±22 | 250±21 | 229±11 | 240±19 | 230±14 | 224±13 | 226±14 | 255±30 | 238±22 | 236±18 |
| VGAT | 276±37 | 301±51 | 293±41 | 270±27 | 301±36 | 271±15 | 338±71 | 272±20 | 293±20 | 291±35 |
| PSD95 | 325±55 | 283±59 | 214±29 | 231±34 | 309±65 | 259±44 | 264±50 | 266±42 | 208±20 | 262±44 |
| GPHN | 257±5 | 252±9 | 243±3 | 260±1 | 280±2 | 270±2 | 267±8 | 258±6 | 260±7 | 261±5 |

**Supplementary Table 7.** Quantification of synaptic puncta density of excitatory and inhibitory markers across layers of human temporal cortex. Measurements (mean ± SEM) normalised to puncta/100 µm^2^.

| Marker/ROI | L1 | L2 | L3 | L4 | L5 | L6 | Mean |
| --- | --- | --- | --- | --- | --- | --- | --- |
| SV2A | 18±1 | 19±1 | 19±1 | 18±1 | 18±1 | 17±1 | 18±1 |
| SYP | 20±5 | 21±2 | 22±2 | 20±4 | 20±3 | 19±3 | 21±3 |
| PSD95 | 41±3 | 44±3 | 42±4 | 40±2 | 36±3 | 32±1 | 39±3 |
| GPHN | 37±8 | 34±4 | 36±3 | 35±5 | 34±9 | 32±8 | 35±6 |

**Supplementary Table 8.** SV2A puncta represent a minor fraction of the total number of synaptic puncta in human temporal cortex layers. SV2A puncta density (top row), summed density of PSD95 and GPHN (middle row), and the percentage of SV2A puncta compared with PSD95 and GPHN (bottom row).

| Marker/ROI | L1 | L2 | L3 | L4 | L5 | L6 | Mean |
| --- | --- | --- | --- | --- | --- | --- | --- |
| SV2A | 18 | 19 | 19 | 18 | 18 | 17 | 18 |
| PSD95 + GPHN | 78 | 78 | 78 | 75 | 71 | 64 | 74 |
| % SV2A | 23 | 25 | 24 | 24 | 25 | 27 | 25 |

**Supplementary Table 9.** Mean intensity of synaptic puncta in layers of the human temporal cortex. Intensity values (a.u.) are shown as mean ± SEM.

| Marker/ROI | L1 | L2 | L3 | L4 | L5 | L6 | Mean |
| --- | --- | --- | --- | --- | --- | --- | --- |
| SV2A | 442±54 | 403±37 | 396±44 | 376±46 | 382±44 | 388±47 | 397±45 |
| SYP | 300±47 | 270±23 | 282±25 | 276±41 | 280±55 | 296±75 | 284±44 |
| PSD95 | 321±29 | 327±42 | 321±19 | 292±16 | 275±23 | 256±16 | 299±24 |
| GPHN | 277±71 | 266±26 | 262±11 | 259±22 | 247±67 | 235±61 | 258±43 |

## Notes

### Competing Interest Statement

The authors have declared no competing interest.

